# Interfacial morphodynamics of proliferating microbial communities

**DOI:** 10.1101/2023.10.23.563665

**Authors:** Alejandro Martínez-Calvo, Carolina Trenado-Yuste, Hyunseok Lee, Jeff Gore, Ned S. Wingreen, Sujit S. Datta

**Affiliations:** Princeton Center for Theoretical Science, Princeton University, Princeton, NJ 08544, USA; Department of Chemical and Biological Engineering, Princeton University, Princeton, NJ 08544, USA; Lewis-Sigler Institute for Integrative Genomics, Princeton University, Princeton, NJ 08544, USA; Physics of Living Systems, Department of Physics, Massachusetts Institute of Technology, Cambridge, MA 02139, USA; Department of Molecular Biology, Princeton University, Princeton, NJ 08544, USA

## Abstract

In microbial communities, various cell types often coexist by occupying distinct spatial domains. What determines the shape of the interface between such domains—which in turn influences the interactions between cells and overall community function? Here, we address this question by developing a continuum model of a 2D spatially-structured microbial community with two distinct cell types. We find that, depending on the balance of the different cell proliferation rates and substrate friction coefficients, the interface between domains is either stable and smooth, or unstable and develops finger-like protrusions. We establish quantitative principles describing when these different interfacial behaviors arise, and find good agreement both with the results of previous experimental reports as well as new experiments performed here. Our work thus helps to provide a biophysical basis for understanding the interfacial morphodynamics of proliferating microbial communities, as well as a broader range of proliferating active systems.

---

Microorganisms do not typically live in isolation; instead, they inhabit spatially-structured multicellular communities, with distinct strains/species occupying distinct spatial domains [1–18]. This spatial structure can have important consequences for the functioning and stability of a community. For example, it can influence how the resident microbes proliferate and interact with each other, resist external stressors, and collectively perform biochemical transformations—with crucial implications for biogeochemistry, the environment, food, health, and industry [5, 19–44]. Hence, considerable research has focused on studying the different morphologies exhibited by microbial communities.

Laboratory studies often focus on communities growing on two-dimensional (2D) planar substrates, both due to their ease of visualization as well as their relevance to the many natural communities that also grow on surfaces. While motility can also influence how microbes spread through their surroundings, in many natural settings microbial colonies predominantly expand through proliferation, which we thus focus on in this work. Under these conditions, it is now well known that a broad array of factors—e.g., differences in competition for nutrients, friction with/adhesion to the underlying substrate, interactions with exogenous or cell-secreted compounds, random fluctuations in proliferation, and mutations [3, 5, 18, 19, 33–39, 42, 45–80]—cause different types of microbes to segregate into monoclonal domains on large scales. But what determines the shape of the *interfaces* between these domains is far less well understood, despite the fact that these interfaces are where the interactions between the different constituents [5, 17] take place. In experiments where two species are grown together, they almost always segregate into bullseye-like patterns, with one species localized to an inner core surrounded by an annular shell of the other. However, the shape of the interface between the two differs from system to system. Sometimes, this interface is smooth (Figure 1i) [60], while in other cases, it is wavy with finger-like protrusions (Fig. 1ii–vi), even though the periphery of the outer shell is smooth [58, 71, 75, 77, 81]. Why these differences in interfacial shape arise has thus far remained a puzzle.

**FIG. 1.**
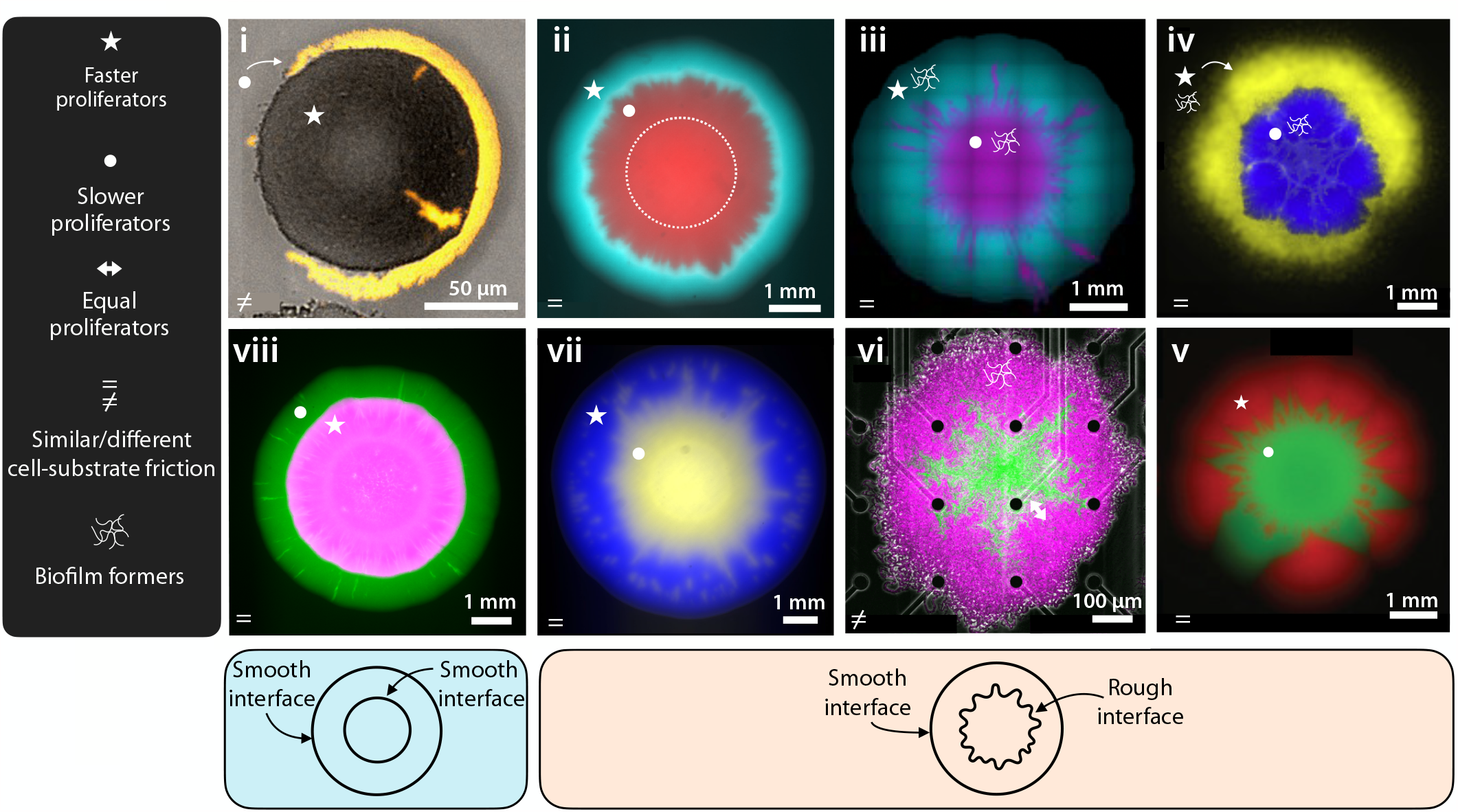
Experimental images of proliferating 2D core/shell-structured microbial communities on flat surfaces. **i**, *Neisseria gonorrhoeae* wild-type (grey inner) and mutant with lower density of type IV pili (yellow outer) [60]. **ii**, Two different strains of *Raoultella planticola* (inner red, outer blue) [77]. **iii**, Two different strains of *Pseudomonas putida* (inner magenta, outer blue) [81]. **iv**, *Pseudomonas aeruginosa* (inner blue) and *Pseudomonas protegens* (outer yellow) [71]; reprinted with permission from Elsevier. **v**, *Saccharomyces cerevisiae* mutant (inner green) and wild-type (outer red) [58]; reprinted with permission from Elsevier. **vi**, Two different strains of *Bacillus subtilis* (inner green, outer magenta) [75]. **vii**, *Pseudomonas rhodeasiae* (inner yellow) and *Raoultella planticola* (outer blue), newly studied in this work. **viii**, *Raoultella planticola* (inner magenta) and *Pantoea agglomerans* (outer green), newly studied in this work. As shown in the left-hand legend, stars/circles indicate faster/slower proliferators, respectively; equal/unequal sign in the lower left of each panel indicates similar/different cell-substrate friction between cell types; additional polymer network symbol indicates cells that produce extracellular polymeric substances. As shown in the legends below, the inner and outer interfaces are both smooth in **i** and **viii**, whereas the inner interface is wavy and unstable in the other panels.

Here, we establish a biophysical description of multispecies/strain microbial communities that provides a foundation to help resolve this puzzle. As a first step, we develop a minimal continuum model in 2D that incorporates two essential features of microbes: proliferation and friction with/adhesion to (hereafter referred to as “friction” for brevity) the underlying substrate. Specifically, we consider a community with two different cell types segregated into a core and a shell, each with distinct proliferation rates and cell-substrate friction. Our theoretical analysis and numerical simulations reveal that the interface between these two domains becomes morphologically unstable—exhibiting wavy, finger-like protrusions as seen in some experiments—when cells in the outer shell proliferate faster or have stronger substrate friction than those in the inner core; otherwise, the interface remains smooth. Moreover, our analysis yields quantitative principles describing when this interfacial instability arises, which we confirm experimentally. Altogether, our work elucidates simple quantitative rules that can help describe the interfacial morphodynamics of microbial communities, and potentially other forms of proliferating active matter [82–106].

## Results

### Model for a proliferating microbial community with two different cell types

Motivated by the experimental observations shown in Fig. 1, we consider a 2D continuum system of two different microbial cell types. Our aim is not to unravel how an initially well-mixed community spatially segregates, but rather to study the morphodynamics of the interface between the different cell types after they have segregated. Thus, our starting point is a community that has already segregated into concentric core-shell domains of differing cell types, as schematized in Fig. 2a. For simplicity, we assume that cells in both domains are effectively incompressible and close-packed at a uniform density *ρ*. Moreover, to decouple proliferation from possible spatial variations in the abundance of nutrients, we assume that the community is under nutrient-replete conditions. Therefore, the cells in each domain proliferate uniformly through space at a constant maximal growth rate *g*_i_, where i = {in, out} refers to the inner core and outer shell domains, respectively. However, as shown in Figs. S5 and S6, we find similar results to those presented in the main text when these assumptions of constant density and proliferation rate are relaxed.

**FIG. 2.**
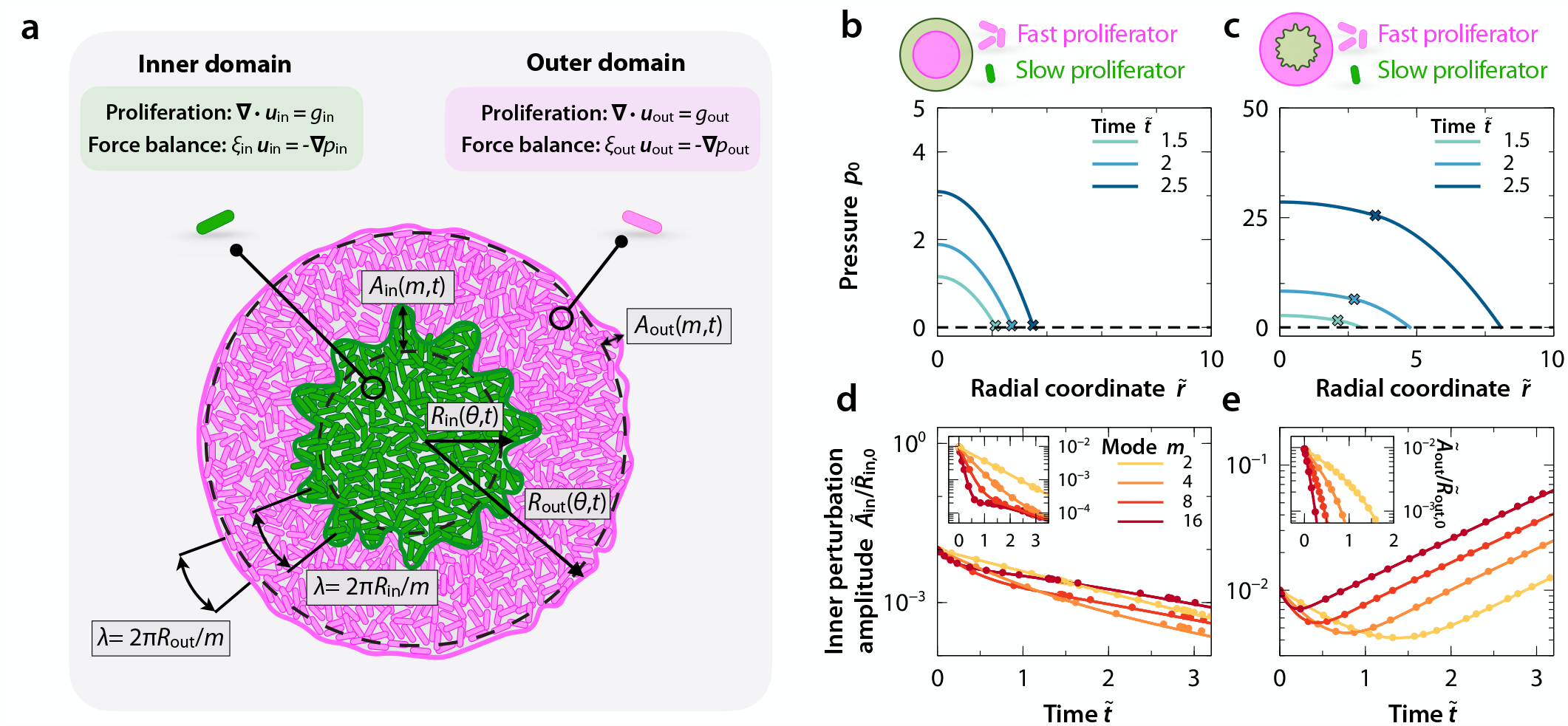
Mathematical model of a proliferating 2D microbial community with two distinct cell types. **a**, Schematic of the 2D continuum system showing an inner domain with one cell type (green) and an outer domain with a different cell type (magenta). 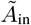 and 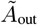 are the time 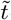-dependent amplitudes of shape perturbations with mode number *m* to the inner and outer interfaces, respectively; 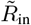and 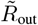are the respective azimuthal angle *θ*-dependent radii of each domain. **b, c**, Base 1D pressure field 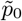 as a function of radial coordinate 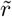 at different times for ℛ = 1.05, *χ* = 1 and **b**,𝒢 = 0.25 (inner domain proliferates faster), **c**, 𝒢 = 2.5 (inner domain proliferates slower). Crosses indicate the position of the interface between the domains. **d, e** Normalized amplitude of perturbations to the inner interface as a function of time 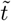 for the same parameter values as in **b** and **c**, respectively; the case of **d** is stable, with perturbation amplitudes that decay over time, whereas the case of **e** is unstable, with perturbation amplitudes that grow with time. The insets to **d** and **e** show similar data, but for the outer interface, indicating that it is always stable. Dots are obtained from full numerical simulations of Eqs. (1)–(2) whereas curves are obtained from linear-stability analysis [Eq. (5)], showing excellent agreement between the two. Variables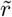and 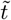in **b**–**e** are normalized by *R*_in,init_ and 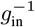, respectively.

The expansion of each domain is caused by cells growing, pushing each other, and proliferating, which can be modeled at the continuum scale as pressure-driven expansion of an “active” fluid [38, 39, 82–84, 91, 107–118]. Hence, we describe the community in terms of a pressure field *p*(***r***, *t*) that is generated by cellular growth and proliferation, and a velocity field ***u***(***r***, *t*) that is proportional to the local pressure gradient ∇*p*; here, ***r*** and *t* denote position and time, respectively. The governing equations for each of the two domains are

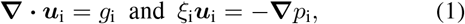

where 0 ≤ *r≤ R*_in_ and *R*_in_ ≤ *r≤ R*_out_ describe the inner and outer domains of radius *R*_in_(*θ, t*) and *R*_out_(*θ, t*), respectively, and *θ* indicates the azimuthal angle (Fig. 2a). Here, *ξ*_i_ is the friction coefficient between cells in domain i and the underlying substrate; *ξ*_i_ relates the pressure gradient driving expansion to the expansion speed, and is taken to be spatially uniform in each domain. [38, 119]. As boundary conditions, we impose continuity of pressure and velocity at the inner interface *R*_in_(*θ, t*), zero pressure at the outer interface *R*_out_(*θ, t*), and a kinematic condition specifying that the velocity of interface i is equal to the velocity of the fluid at that interface:

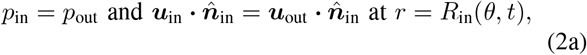

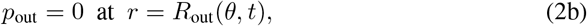

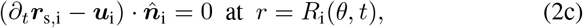

where 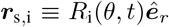 and 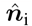 are the vectors describing the position of and unit normal to interface i, respectively, and ***ê***_*r*_ and ***ê***_*θ*_ are the radial and azimuthal unit vectors.

### Dimensionless parameters governing our system

Before solving Eqs. (1)–(2) describing the interfacial morphodynamics, we nondimensionalize these governing equations (Supplemental Materials). To do so, we choose the inverse of the maximal proliferation rate of the inner domain 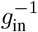 as the characteristic time scale, the initial position of the inner interface *R*_in,init_ as the characteristic length scale, *g*_in_*R*_in,init_ as the corresponding characteristic velocity scale, and 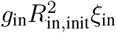as the corresponding characteristic pressure. For ease of notation, all variables presented hereafter are nondimensionalized by these quantities, as indicated by tildes. Nondimensionalizing Eqs. (1)–(2) then reveals three key dimensionless parameters:

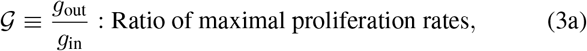

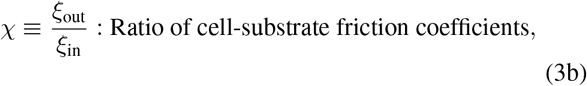

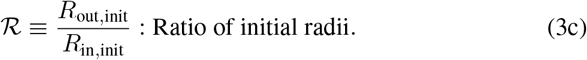

The first two parameters describe differences between the proliferation rates and cell-substrate friction coefficients of the inner core and outer shell domains. The last parameter measures the initial thickness of the outer shell domain. For the results presented in the main text below, we focus on the regime ℛ ≃ 1 often observed in experiments, in which the outer domain has just started to expand. Nonetheless, as shown in Figs. S2 and S3, we find qualitatively similar results when considering larger values of ℛ.

### Base solution of our model

Having developed this model of a proliferating two-domain microbial community, we next obtain a one-dimensional (1D) base solution of Eqs. (1)-(2), denoted by the subscript _0_. To do so, we enforce that the inner and outer interfaces are circularly symmetric. Thus, velocity and pressure depend on the radial coordinate only, 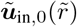 and 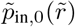, and the positions of the expanding interfaces are only functions of time, i.e., 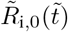. The corresponding 1D solution describing the velocity, pressure, and interfacial positions is

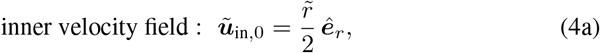

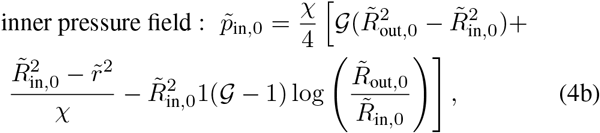

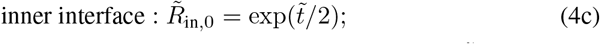

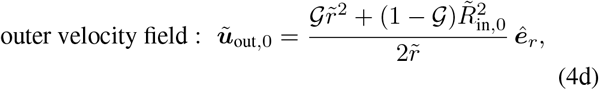

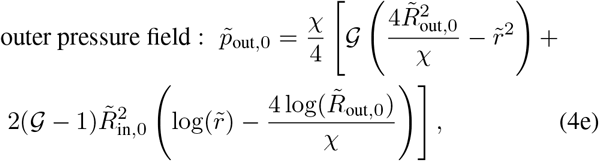

outer interface :

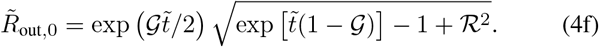

The pressure gradient driving expansion varies throughout the community, and depends on the ratio of proliferation rates 𝒢. Two examples are given by Figs. 2b and c, which show the time evolution of the radial pressure profile 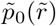 when 𝒢 = 0.25 or 𝒢 = 2.5 with *χ* = 1, corresponding to the opposing cases of more rapid proliferation of the inner or outer domains, respectively. In the former case, the pressure decreases radially outward through the inner domain due to its faster proliferation, and thus is weak in the outer domain. Conversely, in the latter case, the pressure remains high through the entire inner domain due to the faster proliferation around it, decreasing gradually through the outer domain instead. This behavior reflects a fundamental difference between our model and that of an externally driven, passive non-proliferating fluid (*g*_i_ → 0), which corresponds to the classic Saffman-Taylor problem that is well known to have an unstable interface [120–127]. In that problem, the base solution has a uniform velocity and a pressure field that remains linear over time, in stark contrast to our model. Given this difference, we next ask: How do the nonuniform pressure gradients and velocity fields arising from microbial proliferation influence the stability of the inner and outer interfaces in our model?

### Linear-stability analysis reveals that the balance of proliferation rates helps determine interfacial stability

Having obtained a 1D description of the expansion of both inner and outer domains, we next examine the stability of their interfaces. To this end, we perturb Eqs. (1)–(2) with small-amplitude azimuthal disturbances that can be decomposed into Fourier-like modes: 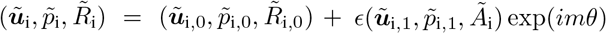, where *Ã*_i_ is the time-dependent amplitude of a perturbation with azimuthal mode number *m*. Introducing this normal-mode decomposition into Eqs. (1) allows us to obtain 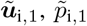, and *Ã*_i_ using the boundary conditions given in Eqs. (2). Since the time scale of domain expansion is of the same order as that describing the growth of perturbations, a standard dispersion relation between the perturbation growth rate and *m* cannot be obtained; the growth or decay of perturbations does not necessarily vary exponentially in time. Instead, interfacial stability is determined by the perturbation amplitudes *Ã*_i_, which are obtained by solving the coupled system of kinematic equations Eq. (2c) at leading order in *ϵ*,

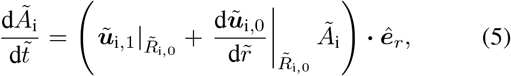

with initial conditions 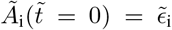. This coupled system of equations does not admit an analytical solution; we therefore solve it numerically and examine the time dependence of the normalized perturbation amplitudes 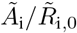 If 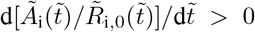, shape perturbations at interface i will grow over time, leading to undulations and the formation of finger-like protrusions, while if 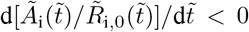, shape perturbations will smooth out over time, yielding a stable, circular interface.

We first explore the two cases shown in Figs. 2b and c, which correspond to *χ* = 1 and 𝒢 = 0.25 or 2.5, respectively. In the former case, both 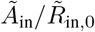 and 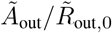 decay monotonically over time for all mode numbers *m*, as shown by the curves in Fig. 2d (main and inset, respectively). Thus, the more rapid proliferation in the inner domain causes both interfaces—between the inner core and outer shell, and at the periphery of the outer shell—to be stable to shape perturbations, smoothing out disturbances and resulting in circular expansion. We confirm this finding using full 2D numerical simulations of Eqs. (1)–(2), with the initial shape of each interface perturbed by the corresponding mode number *m*. The results are shown by the symbols in Fig. 2d, in excellent agreement with the linear-stability analysis.

We observe dramatically different behavior for the latter case of 𝒢= 2.5, corresponding to more rapid proliferation in the outer domain. While 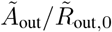 again decays over time for all mode numbers *m* (Fig. 2e, inset), by contrast in this case, 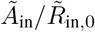 *increases* with time (Fig. 2e, main). This striking result from linear-stability analysis is again confirmed by numerical simulations, as shown by the excellent agreement between the symbols and the curves. Thus, while the periphery of the outer shell is stable to shape perturbations, the interface between the inner and outer domains is not—as observed in corresponding experiments (Fig. 1). Moreover, perturbations with larger mode numbers *m* begin growing sooner than those with smaller mode numbers (different colors in Fig. 2e); that is, short-wavelength perturbations are destabilized faster. The consequences of this interfacial instability are shown by the simulation snapshots in Figs. 3a– c: while the periphery of the outer domain remains stable, the interface between the inner and outer domains quickly becomes unstable, forming wavy, finger-like protrusions similar to those seen in the experiments. This behavior is strikingly different from the classic Saffman-Taylor problem, which requires a difference in resistance across an interface (i.e., *χ* ≠ 1) for it to become unstable [120–127]; here, in stark contrast, the interface is unstable even though *χ* = 1. Altogether, these theoretical and numerical results reveal that the shape of the interface between two different cell types in a microbial community can remain stable, or become destabilized, depending on the ratio of the proliferation rates between the two. Performing the same simulations, but for an interface between slab-shaped—versus concentric circular—domains yields similar results (Figs. S1 and S3), indicating that this phenomenon is not specific to the circular geometry.

**FIG. 3.**
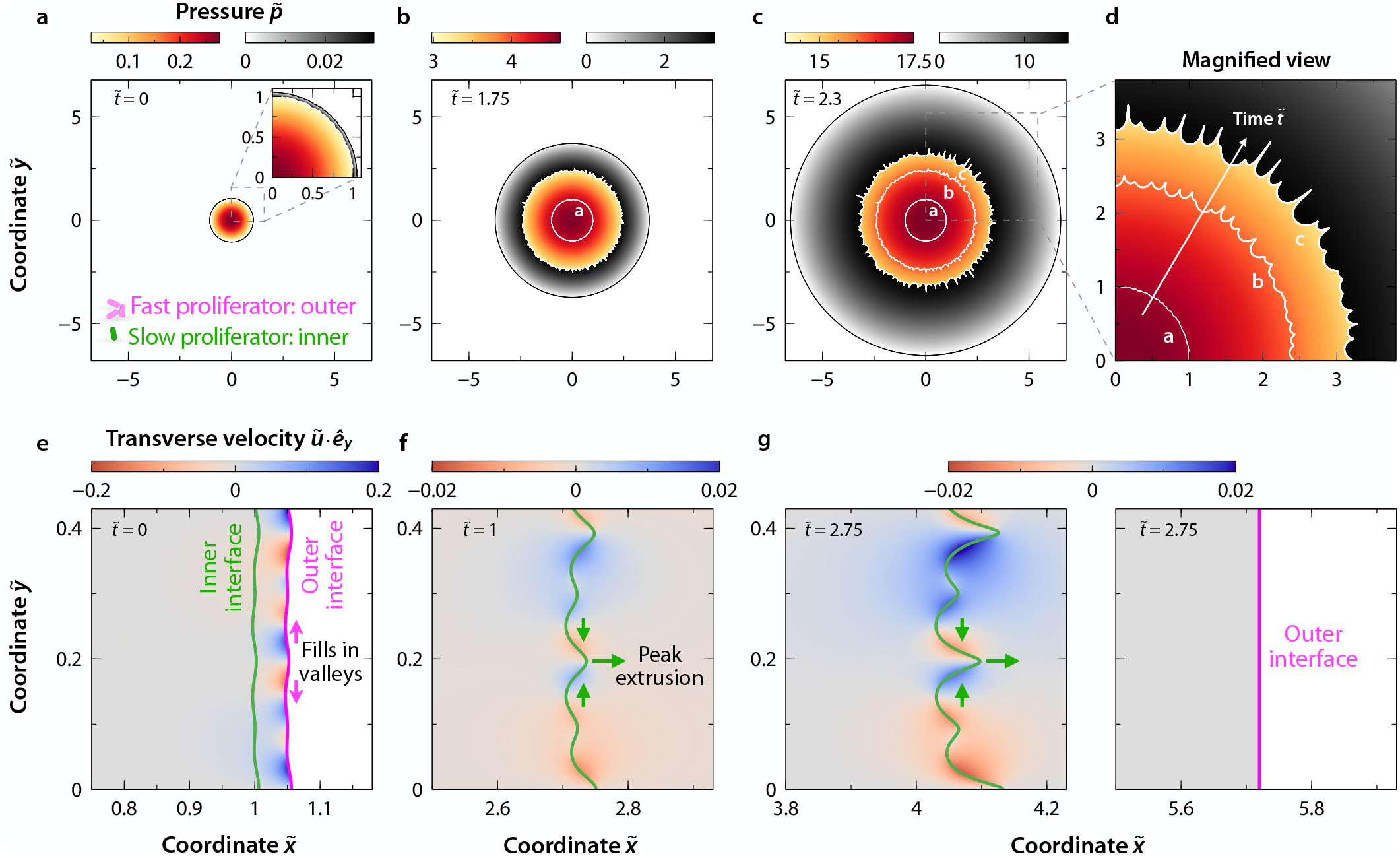
Numerical simulations reveal that differential proliferation transverse to the expansion direction determines interfacial stability. **a**–**d**, Color plots of the pressure field 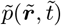 in the inner and outer domains for 𝒢 = 2.5 (outer domain proliferates faster), *χ* = 1, and *ℛ* = 1.05, at **a**, 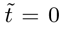, **b**, 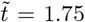, and **c, d**, 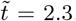. Panel **d** shows a magnified view of **c**, where white curves indicate the shape of the inner interface at different times. The initial shape of both interfaces is perturbed with random white noise. **e**–**g**, Color plots of the transverse velocity 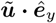 in the inner and outer domains for a slab-shaped community with the same values of 𝒢and *χ* as in **a**–**d**, and *ℛ*= 1.05 (where here *ℛ* ≡ *X*_right,init_*/X*_left,init_; see Supplemental Materials), at **e**, 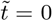, **f**, 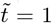 and **g**, 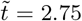. Transverse proliferation in the outer domain squeezes and extrudes the peaks of shape perturbations, destabilizing the interface. In all panels, variables 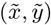 and time 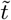 are normalized by *R*_in,init_ and 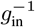, respectively; velocity and pressure are then normalized by *g*_in_*R*_in,init_ and 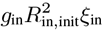, respectively.

### Proliferation transverse to the expansion direction determines interfacial stability

Why is the outer domain always stable? And why does the stability of the interface between the inner and outer domains depend on the ratio of their proliferation rates? Inspecting the profiles of the velocity transverse to the expansion direction in both domains helps to answer these questions. For clarity of visualization, we use simulations of a slab-shaped community with *𝒢* = 2.5, shown in Fig. 3e–g.

First, we focus on the outer interface, shown by the magenta line in Fig. 3e,g. The outer domain proliferates freely, expanding into empty space without any external resistance; indeed, this is the case for any value of 𝒢. Transverse proliferation from the “peaks” of shape perturbations can then fill in the “valleys”, smoothing out these perturbations and stabilizing the outer interface [128]—as quantified by the transverse velocities in Fig. 3e.

We next focus on the interface between inner and outer domains, shown by the green line in Fig. 3e–g. In principle, transverse proliferation in the inner domain can again stabilize shape perturbations of this interface—which indeed is the case when *𝒢 <* 1. However, when the outer domain proliferates faster, 𝒢*>* 1, transverse proliferation in the outer domain has a stronger, opposite effect: it squeezes and extrudes the peaks of shape perturbations, as shown by the green arrows in Fig. 3f–g, destabilizing the interface.

### A morphological state diagram unifies the influence of proliferation and friction in determining interfacial stability

Thus far, we have focused on the influence of differences in the proliferation rate *g*_i_ on interfacial morphodynamics, for inner and outer domains with identical cell-substrate friction (*χ* = 1). However, the different cell types may also differ in their friction coefficient *ξ*_i_ due to, e.g., differences in adhesin expression or production of extracellular polymeric substances (EPS). Hence, we next perform linear-stability analysis over a broad range of 𝒢 and *χ*.

Our results are summarized by the state diagram in Fig. 4, which shows the stability of the interface between inner and outer domains as a function of 𝒢and *χ*, based on the long-time growth rate of perturbations of different mode numbers *m*. The corresponding pressure profiles of the base solution are shown in Fig. S4. The teal shading in Fig. 4 indicates the region of (𝒢, *χ*) parameter space in which both inner and outer interfaces remain stable for all *m*, as in Fig. 2, due to transverse proliferation from peaks of shape perturbations filling in valleys as shown in Fig. 3e. Conversely, the dark orange shading indicates the region in which the outer interface remains stable, but the inner interface becomes unstable, for all *m*, as in Figs. 2e and 3. In this case, the increased friction in the outer domain (*χ >* 1) drives squeezing and extrusion of shape perturbations of the inner interface as in Fig. 3.

**FIG. 4.**
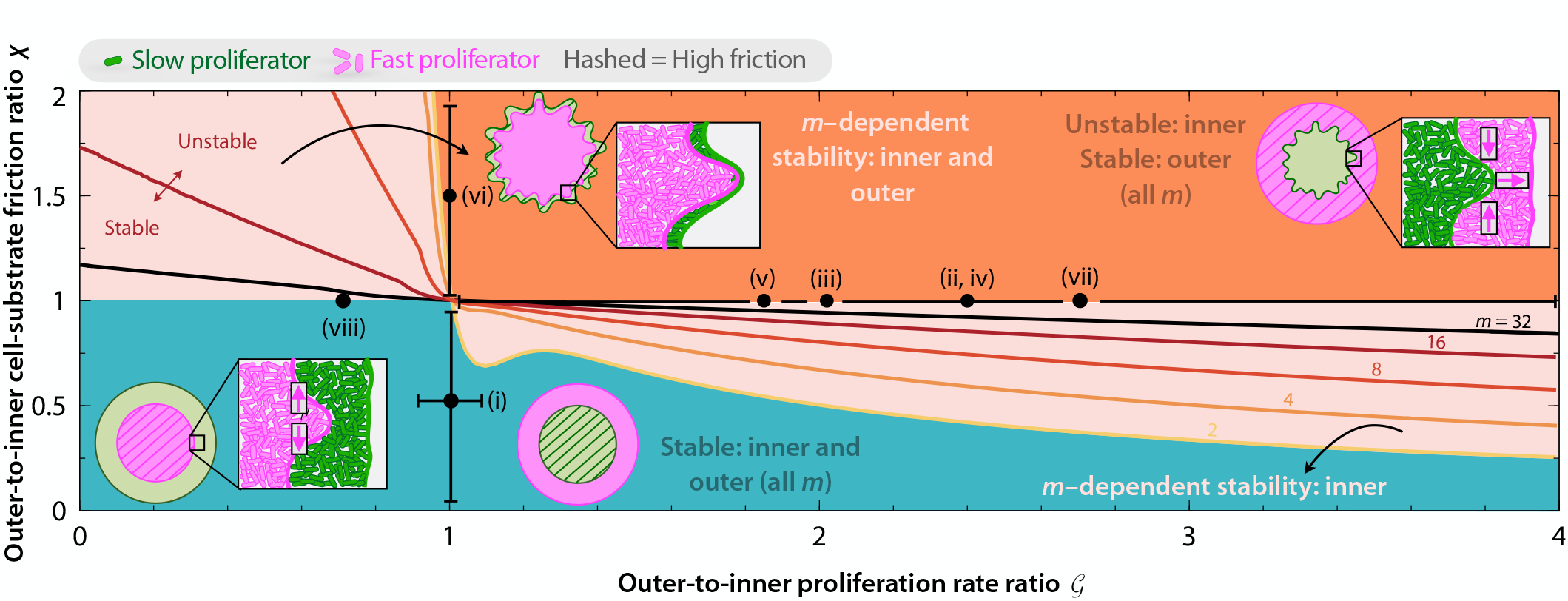
State diagram describes how the balance of differential proliferation and friction determines interfacial stability, in agreement with experiments. Colors show predicted interfacial stability, based on the long-time growth rate of perturbations of different mode number *m*, as a function of 𝒢 and *χ* for a community with ℛ = 1.05. Curves indicate the stable-unstable transition for different *m* modes. Points denote the experiments in Fig. 1; horizontal error bar reflects the reported uncertainty in proliferation rates, while vertical large errors bars indicate that we are unable to quantify the exact value of *χ* for experiments (i) and (vi) and simply know that *χ <* 1 and *χ >* 1, respectively.

The light orange shading in the lower right (𝒢*>* 1, *χ <* 1) indicates an intriguing region in which the outer interface always remains stable, but the inner interface exhibits *m*-dependent stability, with each solid curve indicating the stability boundary for a given mode *m*; above/right of each such curve, the inner interface is unstable to that mode. In this case, the increased friction experienced by the inner domain helps to compensate for the squeezing and extrusion of perturbations at the inner interface induced by more rapid proliferation in the outer domain. The corresponding light orange shading in the upper left (𝒢 *<* 1, *χ >* 1) indicates another intriguing region in which both the outer and inner interfaces exhibit *m*-dependent stability; above/right of each such curve, both interfaces are unstable to that mode *m*. In this case, the stabilizing sideways proliferation of the inner domain is hindered by the increased friction in the outer domain. Moreover, peaks of perturbations at the inner interface proliferate faster than valleys, ultimately pushing the outer interface to also become unstable.

### Theoretical predictions recapitulate experimental observations

The state diagram in Fig. 4 quantifies our central finding: that depending on the proliferation rate and friction ratios 𝒢 and *χ*, the inner and outer interfaces of a two-domain microbial community can be stable, remaining circular, or unstable, generating finger-like protrusions that continue to grow. To what extent do the predictions of our theoretical model capture experimental observations? To address this question, we examine experimental reports in prior literature; details of the analysis of each are provided in the Supplemental Materials.

Fig. 1i, taken from Ref. [60], shows a community of *Neisseria gonorrhoeae* where both inner and outer interfaces are stable. Here, the outer domain is comprised of mutant cells with a smaller density of Type IV pili, thereby reducing cell-substrate friction, compared to the wild-type cells in the interior domain; that is, 𝒢 ≃ 1 and *χ <* 1. As shown in this region of Fig. 4, our theory predicts that both inner and outer interfaces are indeed stable, as observed experimentally.

Figs. 1ii–v, taken from Refs. [58, 71, 77, 81], show communities of bacteria (ii–iv) or yeast (v) where the outer interface remains stable, but the inner interface is unstable. These experiments did not report any differences between the inner and outer cell types with respect to cell-substrate friction (*χ ≃* 1). Rather, in all these experiments, the outer cells proliferated faster than those in the inner domain; that is, 𝒢 *>* 1. As shown by the points in Fig. 4, our theory predicts that the outer interface is stable, but the inner interface is unstable, as observed in the experiments.

Fig. 1vi, taken from Ref. [75], also shows a community of *Bacillus subtilis* where the outer interface remains stable, but the inner interface is unstable. In this case, however, both strains proliferated at a similar rate (𝒢G ≃1). Rather, unlike cells in the inner domain, the outer cells produced extracellular polymeric substances (EPS) that increase cell-substrate friction; that is, *χ >* 1. As shown in this region of Fig. 4, our theory predicts that the outer interface is stable, but the inner interface is unstable, as observed experimentally.

Taken altogether, these analyses show that our theory can rationalize observations made in prior experiments performed on a diverse array of microbial communities. As an additional test of our theoretical predictions, we perform a new experiment with communities of two bacterial species isolated from soil. The outer and inner domains are composed of *Raoultella planticola* and *Pseudomonas rhodeasiae*, respectively, with 𝒢 = 2.7 and *χ ≃* 1 (detailed further in the Supplemental Materials). As shown in Fig. 4, our theory predicts that the outer interface is stable, but the inner interface is unstable—just as we find experimentally, as shown in Fig. 1vii.

None of these experiments investigated the case of 𝒢*<* 1, however. Therefore, as a final test of our theory, we perform another new experiment again with two soil isolates. In this case, the outer and inner domains are composed of *R. planticola* and *P. agglomerans*, respectively, this time with 𝒢 = 0.7 and *χ ≃*1 (Supplemental Materials). As shown in Fig. 4, our theory predicts that both outer and inner interfaces are stable—which our experiment again corroborates, as shown in Fig. 1viii. Thus, the predictions of our theoretical analysis recapitulate experimental observations, both published in previous literature and newly performed in this work, across a broad range of microbial species.

## Discussion

By combining theory, simulation, and experiment, we have established a biophysical description of the morphodynamics of interfaces between domains of two different cell types in a microbial community. For the commonly observed core-shell segregation of two cell types, we found that the balance of the different proliferation rates (𝒢 *≡ g*_out_*/g*_in_) and cell-substrate friction coefficients (*χ ≡ χ*_out_*/χ*_in_) plays a pivotal role in determining interfacial stability. When 𝒢 and *χ* are small, valleys in an interface are transversely filled in by adjacent peaks, resulting in a stable, smooth interface. By contrast, when 𝒢 and *χ* are large, transverse squeezing from valleys causes peaks to become extruded into finger-like protrusions, resulting in an unstable, wavy interface—providing an active matter counterpart to the classic Saffman-Taylor instability exhibited by passive fluid interfaces [120–127]. Notably, our theory quantitatively captures the results of both previously-published experiments, as well as new experiments performed here, across a range of different bacterial and yeast strains [58, 71, 75, 77, 81], as well as in different geometries. We therefore expect that our findings will be generally applicable.

We necessarily made several simplifications and assumptions in formulating our theoretical description. For example, our continuum description neglected orientational ordering of cells, and assumed that the cell proliferation rate *g*, friction coefficient *ξ*, and density *ρ* are constant throughout each distinct domain [129, 130]. There may be nontrivial couplings between the pressure *p* and *g* [131, 132], as well as nontrivial features of cellular rheology, that could change the form of Eq. (1) and potentially give rise to out-of-plane proliferation into the third dimension [53, 133, 134]. Moreover, while our model considers different cell types that interact with each other purely mechanically, they may also regulate each other’s proliferation through other means e.g., via cell-secreted factors such as metabolic byproducts, autoinducers, and toxins [135–138]. Incorporating these complexities into the framework developed here will be a useful direction for future work.

What could the biological implications of the interfacial morphodynamics revealed by our work be? The emergence of wavy fingers at the interface between different cell types could be beneficial for the community. By increasing the length of the interface, this interfacial instability may promote cooperative metabolic interactions between the two domains. Cells located at finger-like protrusions may have better access to any beneficial byproducts released by the other cell type. This instability may also facilitate short-range communication between cells through diffusible signals and metabolites [135–138], or even long-range communication via ion channels [21, 24, 25]. On longer time scales, the increase in interfacial length caused by this instability could also facilitate gene transfer between the different cell types, with possible implications for e.g., the spread of antibiotic resistance [139]. Hence, our work suggests the tantalizing hypothesis that, by regulating cell-scale proliferation/friction, different cell types may be able to regulate large-scale community structure and functioning. Conversely, however, this effect could also be harmful to a community by promoting exposure of the cells to external stressors. Testing these ideas will be an important direction for future research.

By elucidating simple quantitative principles that can help describe the interfacial morphodynamics of microbial communities, our work could also help guide efforts to engineer microbial systems with programmed structures and functions [67, 101–103, 140]. More broadly, our theoretical framework is not restricted to microbes but could be extended to other forms of proliferating active matter, such as developing mammalian tissues [82–99, 105, 106].

## Acknowledgements

A.M.-C. acknowledges support from the Princeton Center for Theoretical Science and the Human Frontier Science Program through the grant LT000035/2021-C. C.T-Y. acknowledges support from the Damon Runyon Cancer Research Foundation through the 2023 Damon Runyon Quantitative Biology Fellowship, the New Jersey Department of Health, the Division of Office of Research Initiatives, and the New Jersey Commission on Cancer Research (NJCCR) through the 2023 NJCCR Postdoctoral Research Grant. H.L. and J.G. acknowledge support from MIT Physics of Living Systems and the Sloan Foundation through grant G-2021-16758. N.S.W. acknowledges support from the NSF through the Center for the Physics of Biological Function PHY-1734030 and the NIH through grant R01 GM082938. S.S.D. acknowledges support from NSF Grants CBET-1941716, DMR-2011750, and EF-2124863, as well as the Camille Dreyfus Teacher-Scholar and Pew Biomedical Scholars Programs, the Eric and Wendy Schmidt Transformative Technology Fund, and the Princeton Catalysis Initiative. We thank Howard A. Stone and Hongbo Zhao for thoughtful discussions.

## Author contributions

A.M.-C. and C.T-Y. carried out the theoretical calculations and numerical simulations, and contributed equally to this work; H.L. carried out the experiments; A.M.-C., C.T-Y., N.S.W., and S.S.D. designed the overall study and analyzed the data; all authors contributed to writing the paper.

## SUPPLEMENTAL MATERIALS

### Dimensionless governing equations

Here we present the dimensionless version of Eqs. (1)–(2), obtained using the characteristic scales described in the Main Text. In particular, we express the system of equations in terms of the proliferation pressure field, which yields:

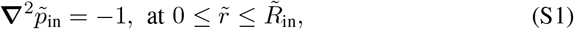

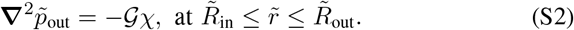

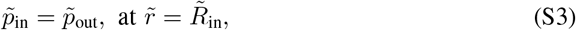

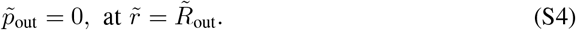

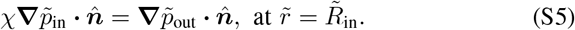

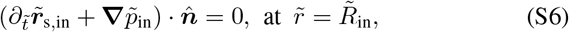

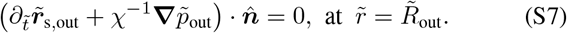

### Slab-shaped communities

Here we investigate the same model as that presented in the Main Text, but for a community that is slab-shaped, as schematized in Fig. S2 rather than circular. To this end, we first obtain a 1D solution of Eqs. (1)–(2) enforcing that the expanding interfaces of both proliferating domains are purely planar, i.e. all the variables depend on the 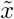 coordinate only and the position of both interfaces are functions of time 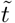 only. The 1D solution reads,

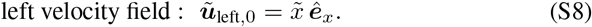

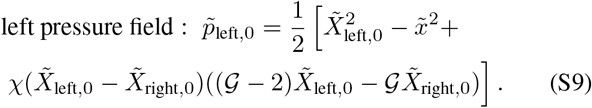

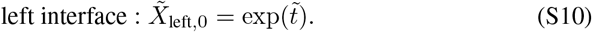

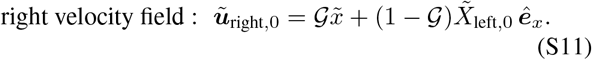

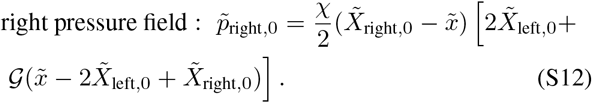

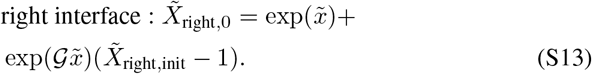

To perform a linear-stability analysis of the above 1D solution, we perturb Eqs. (1)–(2) by small-amplitude disturbances, which are decomposed into Fourier-like modes, i.e. 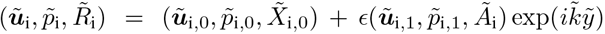, where 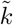 is the wavenumber of perturbations along the 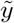 direction, 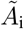 denotes the time-dependent perturbation amplitude of the expanding interfaces, and i = {left, right} . As described in the Main Text for the circular case, we first solve 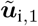 and 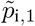 subject to boundary conditions [Eq. (2)], and then 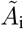 are obtained by solving the kinematic condition at leading order in *ϵ* [Eq. (5)]:

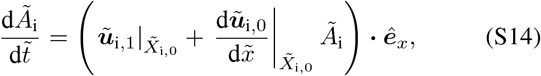

with initial conditions 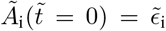. Equivalently to the circular geometry, this coupled system of equations does not admit an analytical solution; we therefore solve it numerically and examine the time dependence of the normalized perturbation amplitudes 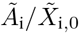. The results of the linear-stability analysis are summarized in Fig. S2, equivalently to Fig. 2 for the circular case, yielding the same results up to metric factors.

### The morphological instability is not affected by nonuniform cell density

Here we consider the case in which cell density *ρ*_i_(***r***, *t*) is not uniform. Hence, the mass and momentum conservation equations read:

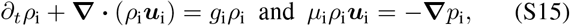

where *μ*_i_ is the cell-substrate friction coefficient per unit density. To solve Eq. (S15), the relationship between the pressure and density fields must be specified. For simplicity, we follow Ref. [141], and consider the simplest possible relation, i.e. *p*_i_(*ρ*_i_) = *G*_i_(*ρ*_i_*/ρ*_i,un_ −1), where *ρ*_i,un_ is an uncompressed density of closely packed cells (i.e. *p*_i_ = 0), and *G*_i_ is a coefficient relating pressure and cell density that reflects the elastic modulus of cells [107]. Equations equivalent to Eq. (S15) but with different pressure-density relationships have been previously employed to model colonies of bacteria [107, 141, 142] and eukaryotic cells [86, 143, 144]. Equation (S15) can be simplified to a reaction-diffusion equation for the density, *∂*_*t*_*ρ*_i_ = (*μ*_i_*ρ*_i,un_)^−1^**∇**^2^*ρ*_i_+*ρ*_i_*g*_i_, where the expanding interfaces move according to ***u***_i_ = −*G*_i_**∇***ρ*_i_*/*(*μ*_i_*ρ*_i,un_*ρ*_i_). Equivalently to the incompressible case in the Main Text, we impose continuity of pressure and velocities, and kinematic conditions at both interfaces, and zero pressure at the outermost interface.

To nondimensionalize the new system of equations, we choose the same characteristic scales for the length, velocity, and time. The characteristic pressure scale is modified to 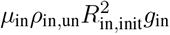. Regarding the density scale, we choose *ρ*_i,un_ for each density field. Using these scales we obtain the following dimensionless system of equations:

**FIG. S1.**
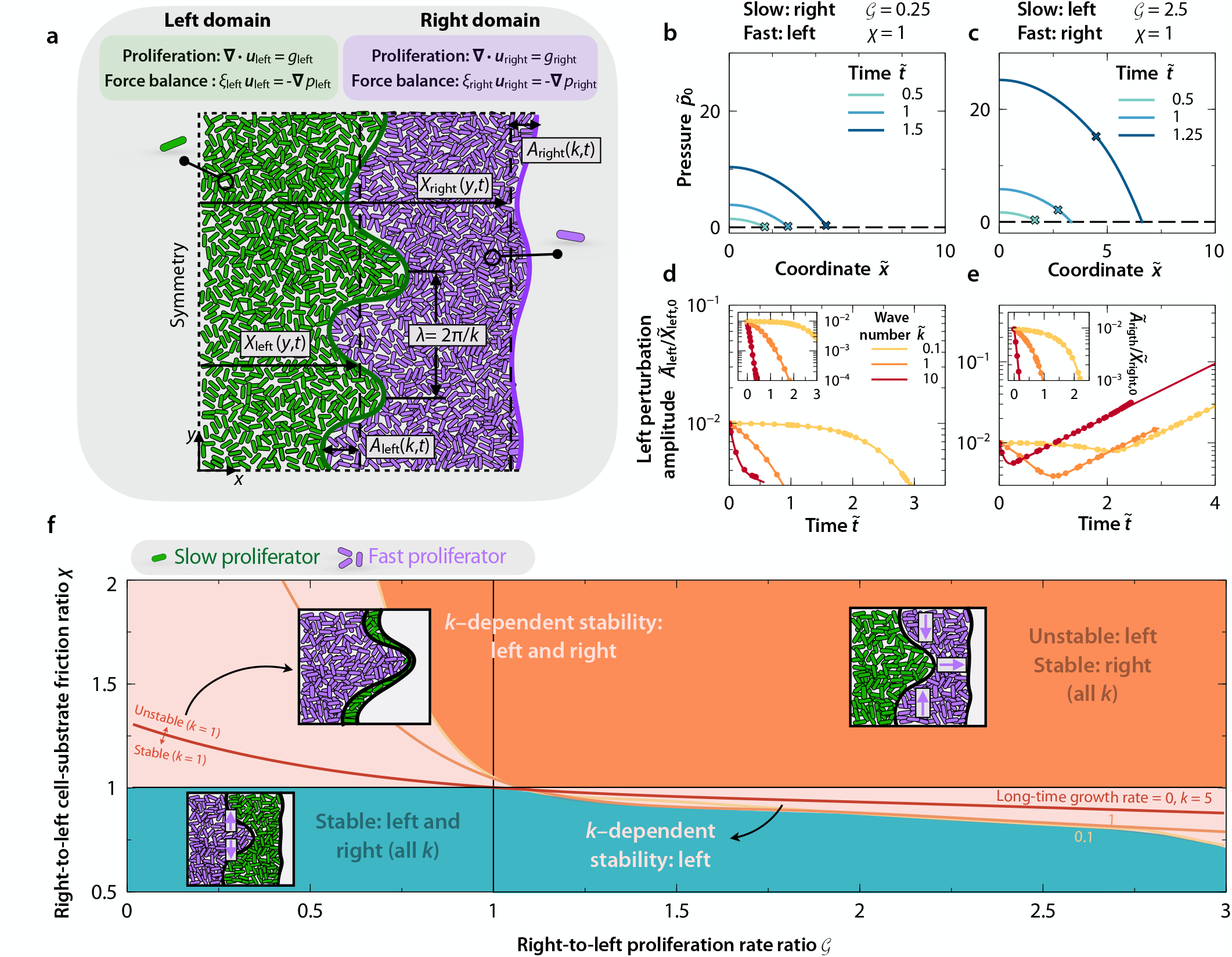
**a**, Schematic of slab-shaped 2D continuum system showing a left domain with one cell type (green) and an right domain with a different cell type (magenta). 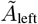 and 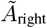 are the time 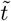-dependent amplitudes of shape perturbations with wavenumber *k* to the inner and outer interfaces, respectively; 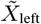 and 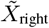 are the respective 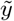-dependent widths of each domain. **b**– **e** are identical to the panels in Fig. 2, but for this community in this new coordinate system. **f**, State diagram identical to Fig. 4, but for this community in this new coordinate system, for perturbations indexed by wavenumber 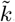. These calculations are for ℛ = 1.05, where here 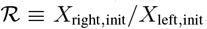. As shown by all these panels, the results for a proliferating microbial community in a slab geometry are similar to those obtained for a circular geometry.

Mass conservation :

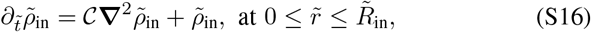

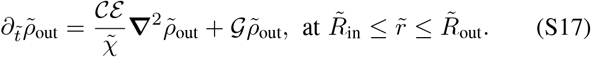

Stress balance:

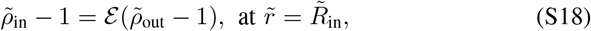

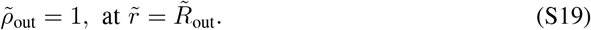

Continuity of velocity:

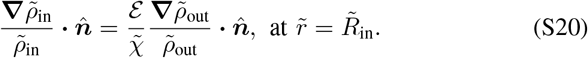

Kinematic condition :

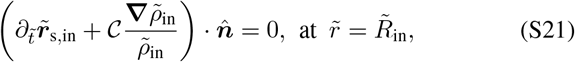

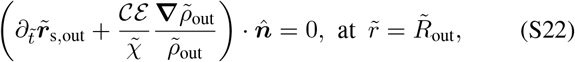

where the new key dimensionless parameters are:

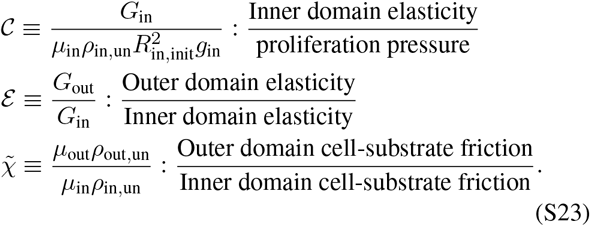

**FIG. S2.**
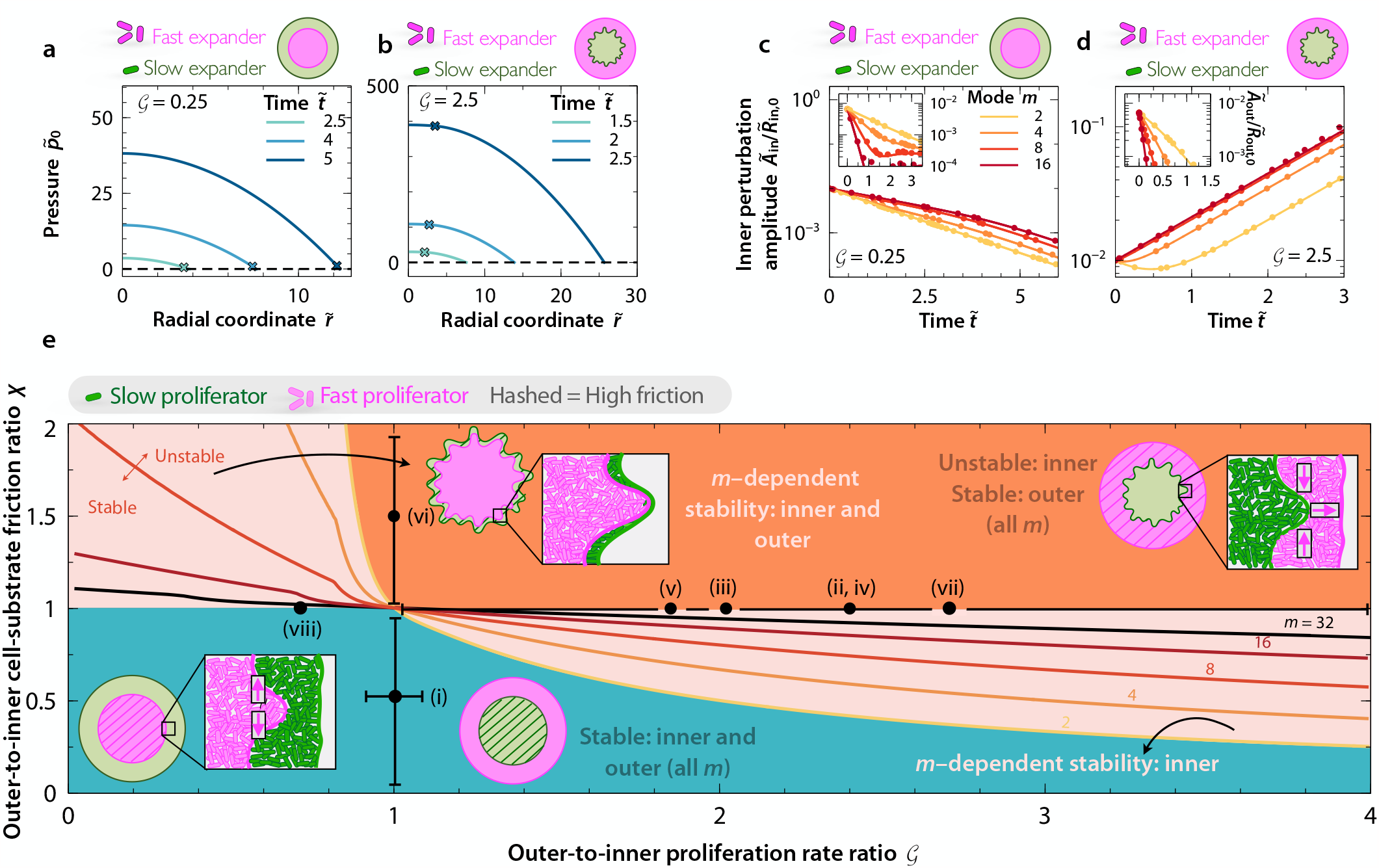
Model predictions for a community with *ℛ* = 1.5 show qualitatively similar results to those for the community with *ℛ* = 1.05 presented in the main text. **a**–**d** are identical to panels **b**–**e** of Fig. 2, but for *ℛ* = 1.5. Similarly, **e** is identical to Fig. 4, but for *ℛ* = 1.5.

To test whether a nonuniform cell density affects the interfacial instability, we perform full 2D numerical simulations of Eqs. (S16) in the circular geometry, imposing an initially uniform density 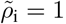. For simplicity, we assume that 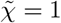 and ℰ = 1, i.e. both domains display the same cell-substrate friction and proliferation pressure-density relation, 𝒞 = 1, 𝒢 = 2.5, and *ℛ*= 1.05 (same as in Fig. 4a–c). Figure S5 displays snapshots of the cell density field 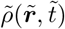 in both domains at times 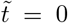 (panel a), 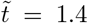 (panel b), and 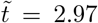 (panel c). As 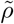 can vary spatially, the expansion is slower than in the incompressible case. Indeed, while the density increases locally with an exponential form at sufficiently long time, the radii of both domains do not grow exponentially. Moreover, at time 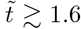, a high-density ring is created at the interface of the contact interface between the two domains (rightmost panels). This high density region travels outwardly with time, expanding the outer domain 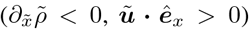 and compressing the inner domain 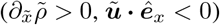. Nonetheless, despite this compression effect whereby the inner interface starts moving inwards, the instability remains unaltered as shown in panels b and c, being identical to that of the incompressible case displayed in Fig. 4a–c. From a biological perspective, the large local increase in cell density shown in Fig. S5 is not realistic. Once the cells are closely packed, the 2D density could not increase as much as we observe in our simulations. Increasing the elastic modulus, considering nutrient-limited growth, and allowing expansion into the third dimension would address this limitation. However, here we focus on this extreme case to demonstrate that even highly compressible cells would still exhibit the same interface morphodynamics.

### Nutrient-limited proliferation does not preclude the morphological instability

To investigate how nutrient-limited proliferation may influence the interfacial instability, we modify Eqs. (1) and consider a nutrient-dependent proliferation rate given by the Monod equation, i.e. **∇** · ***u***_i_ = *g*_i_*c*_i_*/*(*c*_half,i_ + *c*_i_), where *c*(***r***, *t*) denotes the local concentration of nutrients at the inner and outer domains. We assume that nutrient dynamics obeys a standard reaction-diffusion equation that incorporates diffusion and uptake in both domains, i.e. *∂*_*t*_*c*_i_ = *D* **∇** *c*_i_ *k*_i_*ρ*_i_*c*_i_*/*(*c*_half,i_ + *c*_i_), where *k*_i_ is the saturated nutrient uptake rate per cell, *c*_half,i_ is the characteristic constant of the Monod equation, and *D* is the nutrient diffusion coefficient. Moreover, we have assumed that both cellular proliferation and nutrient uptake obey the same Monod dependence. As for the boundary and initial conditions we impose a constant concentration of nutrients *c*_out_ = *c*_0_ at the outer interface ***r*** = ***r***_s,out_, nutrient continuity, i.e. *c*_in_ = *c*_out_ and 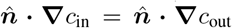 at the inner interface ***r*** = ***r***_s,in_, and *c* = *c*_0_ at *t* = 0. We use *c*_0_ to nondimensionalize the nutrient concentration, which yields the following dimensionless nutrient equation, 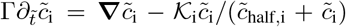, where 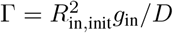 compares cellular proliferation and nutrient diffusion time scales, 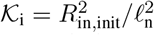, compares the initial radius of the inner domain with the nutrient penetration length, 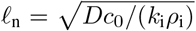, and 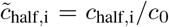 compares the half-velocity constant of each domain with the initial nutrient concentration *c*_0_.

**FIG. S3.**
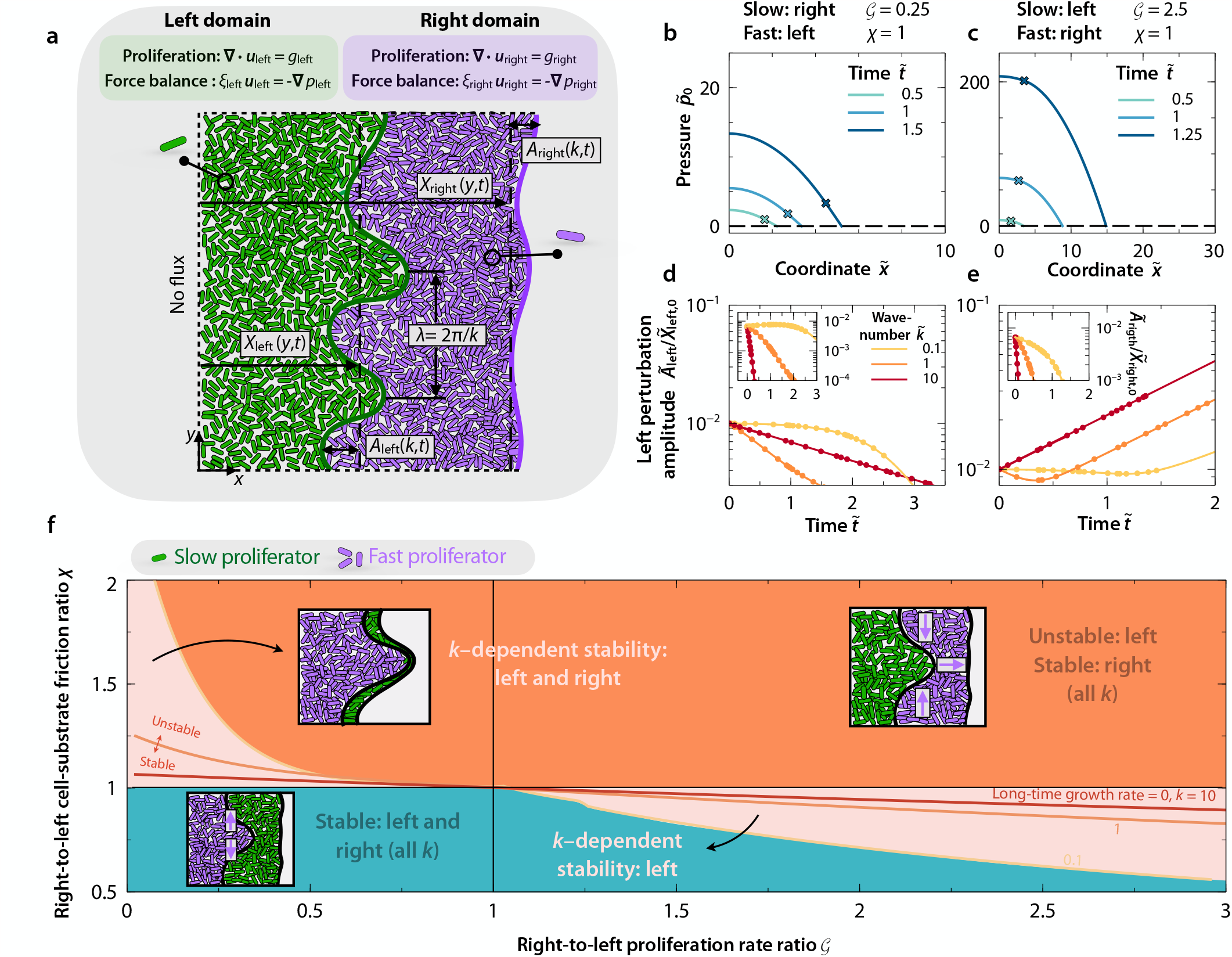
Model predictions for a community with *ℛ*= 1.5 show qualitatively similar results to those for the community with *ℛ* = 1.05 in Fig. S1. **b**–**e** are identical to panels **b**–**e** of Fig. S1, but for *ℛ* = 1.5. Similarly, **f** is identical to Fig. S1f, but for *ℛ* = 1.5.

Figure S6 displays color plots of the local nutrient concentration 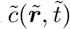 at different times obtained from time-dependent 2D simulations including nutrient diffusion and uptake, using the same values of 𝒢, ℛ, *m*, and *ϵ* as in Fig. 4. Regarding nutrient related dimensionless parameters, we consider Γ ≪ 1 and use 𝒦_in_ = 0.4, 𝒦_out_ = 1, and 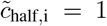 for simplicity. As expected, the expansion of both domains slows down as nutrient becomes depleted in their interior, but we find the same type of morphological stability arising at the expanding interface of the inner domain, independent of the values of the dimensionless parameters related to diffusion and uptake. At sufficiently long time, e.g. 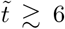 in Fig. S6, the inner domain stops proliferating as it runs out of nutrients, but the rough interface remains without smoothing out, in agreement with recent experiments [77]. Thus, as expected, nutrient-dependent proliferation only modulates the morphological instability of the inner domain by inhibiting proliferation as nutrient becomes depleted, but is not able to stabilize the proliferating interface. Altogether, linear-stability analysis and numerical simulations of our model suggest that such morphological instabilities may be generic, independent of environmental conditions.

**FIG. S4.**
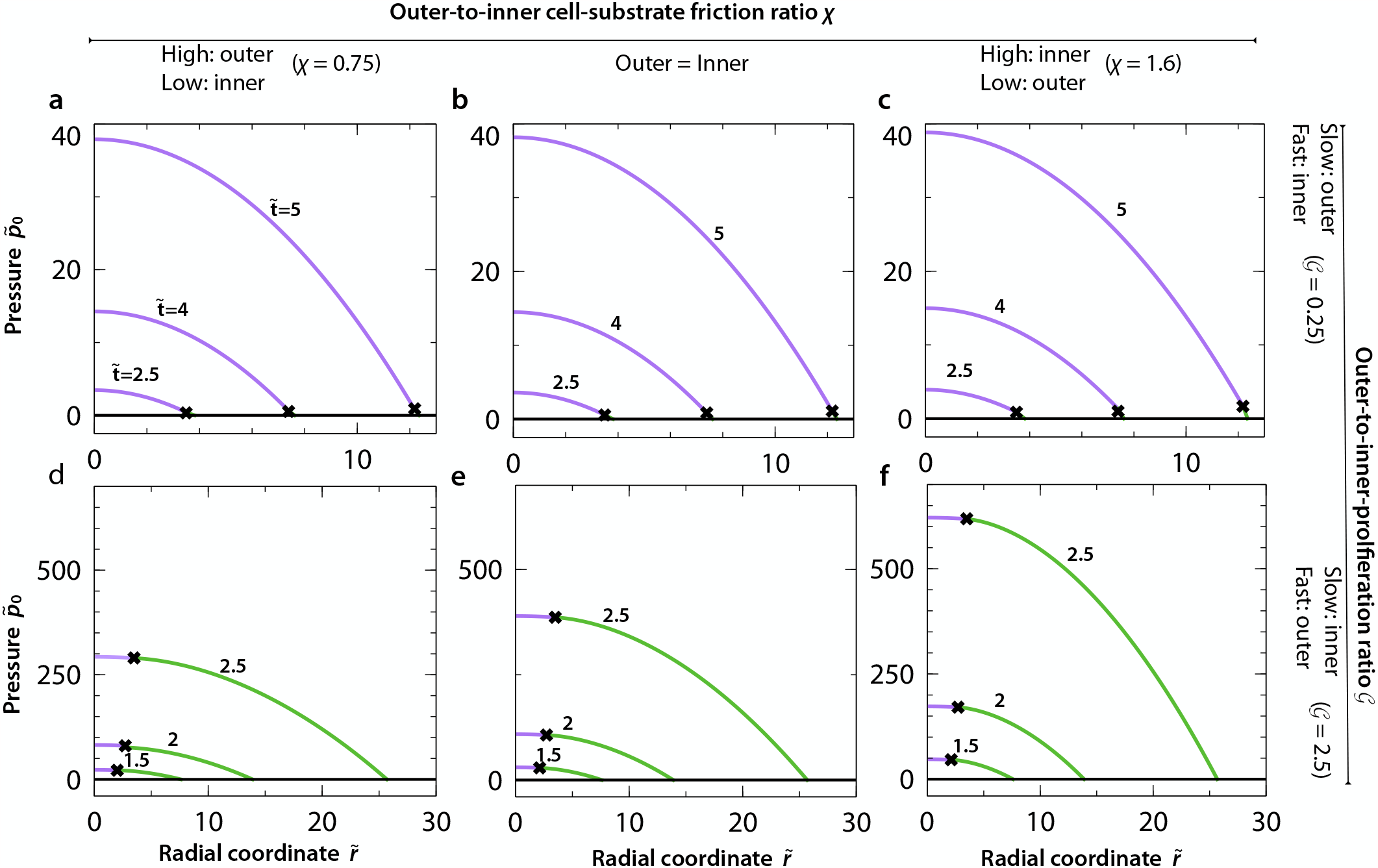
Base 1D pressure field 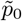 as a function of the radial coordinate *r* at different times for ℛ = 1.05 and different values of 𝒢 and *χ*. Crosses indicate the position of the interface between the domains.

**FIG. S5.**
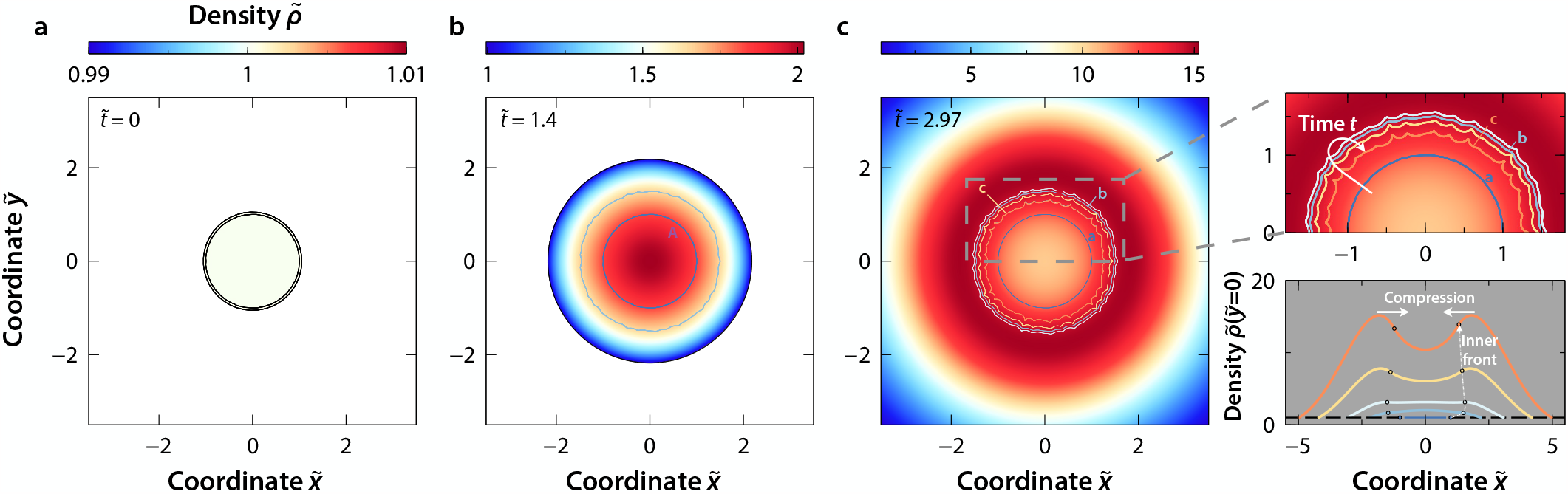
**a**-**c**, Color plots of the density field 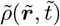 for the same values of 𝒢 as in Fig. 4a–c, and ℛ = 1.05, 𝒞 = 1, ℰ = 1, and 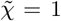, at times 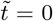, 1.4, and 2.97. The rightmost panels show a zoomed-in view of the density field in **c** (top), and 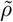 at 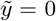 as a function of the 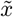 coordinate (bottom), also displaying intermediate times.

### Measurements and estimates of the dimensionless parameters *𝒢* and *χ* for previously published experiments

Here we detail the reported measurements for *g*_i_ of the experiments displayed in Figs. 1. These values are used to compute 𝒢 and determine the position of each experiment on the state diagram shown in Fig. 4. Regarding values of *χ*, as detailed in the Main Text, we consider that *χ* = 1 in all experiments except for Figs. 1i and vi. In the experiment shown in Fig. 1i, the outer domain is a mutant, which exhibits a smaller density of type IV pili and emerges from the inner domain. As discussed by the authors, the outer domain displays weaker cell-substrate friction, as a consequence of reduced pilus density, thus we consider *χ <* 1. In the experiment of Fig. 1vi, the strain in the outer domain is immotile and secretes EPS, while cells in the interior domain are motile and do not secrete EPS, thus we consider *χ >* 1, as presumably the outer domain exhibits a larger friction with the substrate due to EPS-substrate adhesion. We assume *χ* = 1 for the other experiments in Fig. 1, since the two strains are grown on the same substrate, secrete the same EPS in cases of the biofilm formers, and exhibit the same cell shape.

**FIG. S6.**
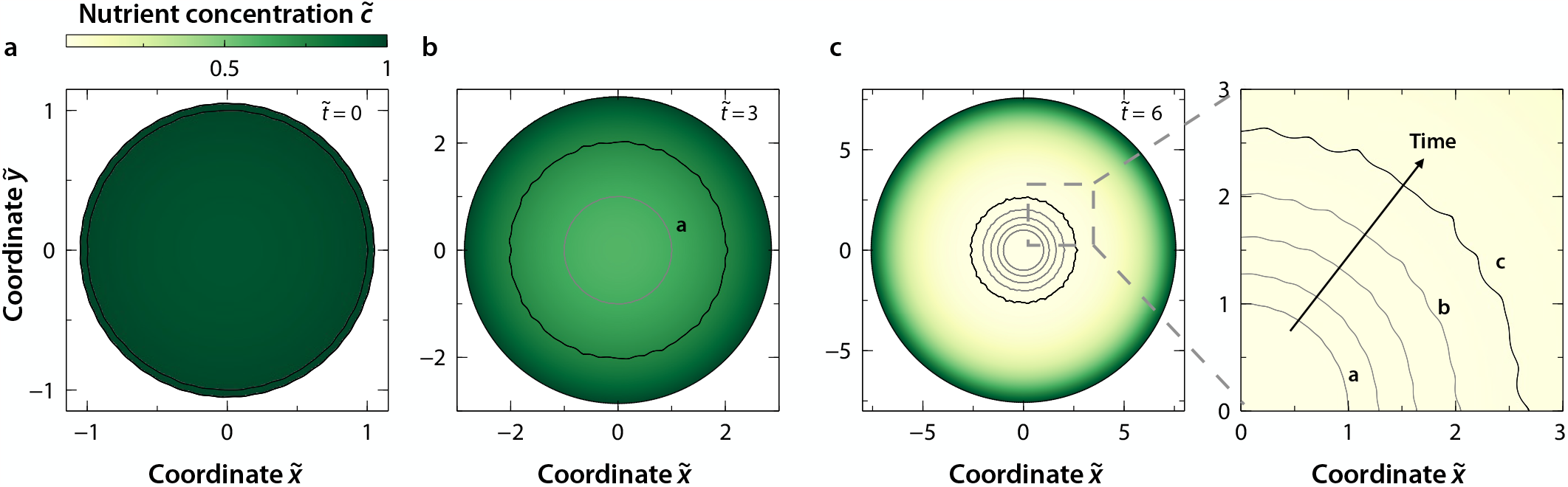
**a**-**c**, Color plots of the 2D nutrient concentration field 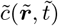 for the same values of *𝒢* as in Fig. 4a–c, *ℛ* = 1.05,Γ ≪ 1, *𝒦*_in_ = 0.4, *𝒦*_out_ = 1, and 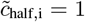, at times 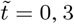, and 6. Grey curves indicate the shape of the inner interface at previous times.

The experiments displayed in Figs. 1ii and 1iv are obtained from Ref. [77] and [71], respectively. In these studies the proliferation rates of the two strains are not reported, but we assume that the outer domain outgrows the inner domain as the former keeps proliferating over time, whereas cells of the inner domain stop proliferating or prolfierate at a much slower rate, similarly to experiments in Ref. [38]. Regarding the experiment shown in Fig. 1v adapted from Ref. [58], *g*_in_ ≃ 0.012 h^−1^, and *g*_out_ ≃ 0.023 h^−1^, thus *𝒢* ≃ 1.9. Experiments displayed by Figs. 1iii are from Ref. [81], in which *g*_in_ ≃ 0.29 h^−1^, and *g*_out_ ≃ 0.58 h^−1^, thus *𝒢* ≃ 2. Finally, *𝒢* ≃ 1 for the experiment shown in Figs. 1vi as reported in Ref. [145].

### New experiments using bacteria isolated from soil

*Raoultella planticola, Pseudomonas rhodeasiae*, and *Pseudomonas agglomerans* strains were isolated from a soil sample (MIT Killian Court, Cambridge, MA) and were tagged with two different fluorescent proteins mScarlet-I (red) and GFP2 (green) by insertion of plasmids pMRE145 and pMRE132 respectively. For growth substrates, we prepared stiff agar plates with 1X Luria-Bertani media (LB, 2.5% w/v; BD Biosciences-US) and 1.5% w/v of agar (BD Bioscience-US). We also added 1X Chloramphenicol (Cm, 15mg/L, prepared from 1000X solution) for constitutive expression of fluorescence. For each agar plate, 4mL of media was pipetted into a petri dish (60X15mm, sterile, with vents; Greiner Bio-one), and was cooled overnight (15 hours) before inoculation.

At the start of the experiment, for each strain, −80°C glycerol stock was streaked on a separate plate and proliferated for 2 days. Then a colony from each strain was picked up and put into a 50mL Falcon Tube filled with 5 mL of liquid media (1X LB and 1X Cm). Bacterial cultures were grown overnight at 30°C under constant shaking 1350 rpm (on Titramax shakers; Heidolph). We then diluted the cultures to total density of optical density 0.1 (OD_600_) using a Varioskan Flash (Thermo Fisher Scientific) plate reader. The OD_600_-standardized cultures were mixed in 1:1 volume fractions, and we then gently placed a droplet of 1.5 *μ*L inoculant at the center of an agar plate. After inoculation, each colony was grown at 30°C for 120 hours. At fixed times after inoculation, each plate was put on a stage of Nikon Eclipse Ti inverted light microscope system. 10X magnification was used for whole-colony images. Fluorescent images were taken using Chroma filter sets ET-dsRed (49005) and ET-CFP (49001) and a Pixis 1024 CCD camera.

**FIG. S7.**
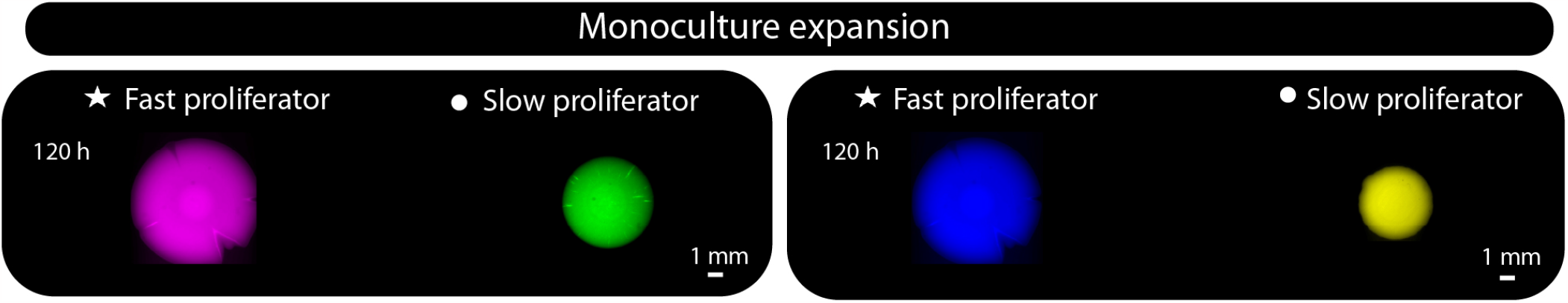
Monoculture expansions for experiments shown in Fig. 1viii (left) and Fig. 1vii (right); the area of each domain was used to determine proliferation rates.

